# A Standardized In Vitro Platform for Senolytic Drug Discovery in Human Musculoskeletal Cells

**DOI:** 10.64898/2026.08.20.746082

**Authors:** Hosni Cherif, Sami Alsabri, Jean A. Ouellet, Lisbet Haglund

## Abstract

Cellular senescence contributes to the progression of many age-related musculoskeletal diseases. Cellular senescence is a biological state that arises from replicative exhaustion and various cellular stressors, including elevated oxidative stress, mitochondrial dysfunction, mechanical overload, and chronic exposure to pro-inflammatory cytokines and proteases. Although senolytic agents show promise for eliminating senescent cells, their translation has been hindered by the lack of physiologically relevant and scalable in vitro screening methods. In the present study, we developed a standardized, physiologically relevant senescence-induction model and validated a metabolic activity assay as a rapid, scalable method for screening senolytic compounds. We used primary human intervertebral disc cells (IVD) as an example, but the workflow applies to many other cell types. To mimic inflammatory and oxidative stress, we used a combination of TLR-2 activation (Pam2CSK4) and tert-butyl hydroperoxide (tBHP), a potent ROS generator. Senescence induction was validated by quantifying β-galactosidase fluorescence intensity, β-gal enzymatic activity, and the expression of the p16 senescence marker across 3 IVD cell types: nucleus pulposus (NP), inner annulus fibrosus (iAF), and outer annulus fibrosus (oAF) cells. The combined Pam2CSK4 + tBHP exposure generated a robust senescent phenotype across all 3 IVD cell types, with oAF cells exhibiting the strongest increases in β-gal fluorescence, β-gal enzymatic activity, and p16 expression. We then used oAF cells to evaluate if the metabolic activity assay (Alamar Blue) could be used to determine both cytotoxicity of senolytic drugs in non-senescent cells and senolytic activity in a mixed population of senescent and non-senescent cells. We validate the method by comparing metabolic activity results with β-gal enzymatic activity and p16 expression in induced and non-induced cells following exposure to three known senolytics (o-Vanillin, RG-7112, and ABT-199). The metabolic activity assay reliably identified a therapeutic window in which the three senolytics were non-toxic to non-senescent cells while selectively reducing metabolic activity in a mixed population of senescent and non-senescent cells. The reductions in metabolic activity in the mixed population correlated with decreases in SA-β-gal enzymatic activity and p16 expression, validating metabolic activity as a sensitive and scalable senolytic readout.

## Introduction

Cellular senescence has emerged as a fundamental biological mechanism contributing to aging and the development of chronic musculoskeletal disorders. Senescent cells accumulate in degenerating tissues and promote disease progression by secreting pro-inflammatory cytokines, proteases, and other components of the senescence-associated secretory phenotype (SASP) ^1-3^. Because of their pathological role, selective elimination of senescent cells using senolytic agents has gained considerable attention as a potential therapeutic strategy for age-related musculoskeletal diseases, including osteoarthritis, osteoporosis, and intervertebral disc (IVD) degeneration. Although numerous compounds have demonstrated promising senolytic activity in preclinical studies, successful translation into clinical applications has remained challenging, highlighting the need for improved and physiologically relevant preclinical screening platforms.

One important limitation in senolytic drug discovery is the availability of in vitro models that accurately reflect the senescent cell populations present in diseased tissues. Despite encouraging preclinical results, clinical translation of senolytics has been limited. Early trials evaluating a combination of Dasatinib and Quercetin in idiopathic pulmonary fibrosis and diabetic kidney disease demonstrated safety but modest efficacy, while senolytic candidates such as UBX0101 failed to show clinical benefit in osteoarthritis ^4, 5^. These findings highlight a critical challenge that many preclinical senolytic assays rely on senescent fibroblasts or immortalized cell lines, which do not recapitulate the biology of senescent cells from a distinct tissue type. Primary musculoskeletal cells isolated from human tissues typically require expansion in culture before experimentation. During this process, proliferative non-senescent cells are preferentially enriched, resulting in cultures that do not accurately represent the in vivo cellular composition of ageing and degenerating tissues. Consequently, experimental induction of senescence is often necessary to generate cell populations suitable for evaluating senotherapeutic compounds. Multiple approaches have been developed to induce senescence in vitro, including replicative exhaustion, genotoxic stress, oxidative stress, irradiation, and inflammatory stimulation ^6-9^. However, the optimal induction strategy is highly dependent on the tissue and cell type under investigation and should ideally reproduce the molecular and environmental stressors driving disease in vivo.

Here, we use intervertebral disc degeneration as a model system for studying cellular senescence in musculoskeletal tissues. Degenerating discs and other musculoskeletal (MSK) tissues are characterized by a chronic inflammatory microenvironment enriched in cytokines such as TNF-α, IL-1β, and IL-6, together with elevated oxidative stress resulting from impaired nutrient transport, mitochondrial dysfunction, and mechanical overload. The inflammatory and oxidative stress promote DNA damage, cellular dysfunction, and activation of senescence pathways, leading to the accumulation of senescent disc cells and progressive tissue degeneration ^10-13^. Previous studies from our group and others have demonstrated the presence of senescent cells in human IVD tissues and have shown that senolytic interventions can selectively target these populations (Cherif et al., 2020; Mannarino et al., 2023). Therefore, reproducing the combined inflammatory and oxidative stress conditions observed in primary MSK cells may provide biologically relevant cell populations for in vitro evaluation of candidate senotherapeutics. In fact, the same senolytic drugs have shown efficacy in isolated cells from these tissues where senescence was induced by inflammatory and oxidative stress as described here.

While establishing a reliable senescence induction model is essential, a second challenge is the identification of screening methods that are sufficiently rapid, inexpensive, and scalable for routine senolytic testing. Conventional senescence assessment relies on markers such as senescence-associated β-galactosidase (SA-β-gal) activity and expression of cell-cycle regulators including p16^INK4a^. Although these markers provide important biological validation, they can be labour-intensive and less suited to high-throughput applications. A simple, rapid, and scalable screening approach capable of identifying compounds with selective senolytic activity, while excluding those that are toxic to non-senescent cells, would facilitate the evaluation and prioritization of larger numbers of candidate molecules before more detailed validation is performed.

In the present study, we developed and validated a reproducible senescence induction protocol combining inflammatory stimulation by activation of Toll-Like-Receptor 2 (TLR-2) with Pam2CSK4 and oxidative stress induced by tert-butyl hydroperoxide (tBHP) in primary human IVD cells. Using established senescence markers, including SA-β-gal activity and p16 expression, we validated the senescence induction resulting from this approach. We subsequently investigated whether a simple metabolic activity assay could serve as a rapid and cost-effective surrogate screening method for senolytic activity. We validated this approach using three well-characterized senolytic compounds, o-Vanillin, RG-7112, and ABT-199, and demonstrated a practical screening strategy that combines physiologically relevant senescence induction with a scalable assay for senolytic drug discovery. By combining physiologically relevant senescence induction with a simple metabolic activity readout, this strategy has the potential to facilitate high-throughput senolytic discovery and accelerate the development of therapies targeting age-related musculoskeletal diseases.

## Results

### 1. Cytotoxicity and selection of tBHP concentration

Studies across different connective tissue cells have shown that TLR activation generates a cell-specific inflammatory environment (Joosten et al., 2016; Miller et al., 2019). Here we used Toll-Like Receptor-2 (TLR-2) activation (Pam2CSK4) at the previously validated concentration of 100 ng/ml to induce the synthesis of cell-type-specific inflammatory mediators (Mannarino et al., 2021; Krock et al., 2016, 2017), while oxidative stress was induced by Tert-butyl hydroperoxide (tBHP). (Park et al, 2002). To determine a non-toxic and biologically relevant concentration range, we first determined the cytotoxicity of tBHP alone and then combined with Pam2CSK4 (100 ng/mL). No significant change in metabolic activity was observed at the tHBP concentrations tested (**Figure 1A**). We confirmed the results by evaluating the cell viability across the three IVD cell populations, nucleus pulposus (NP), inner annulus fibrosus (iAF), and outer annulus fibrosus (oAF), using the Live/Dead assay. Both tBHP and Pam2CSK4+ tBHP maintained metabolic activity and cell viability at the tested concentrations across the IVD cell populations (**Figure 1B**).

**Figure 1.**
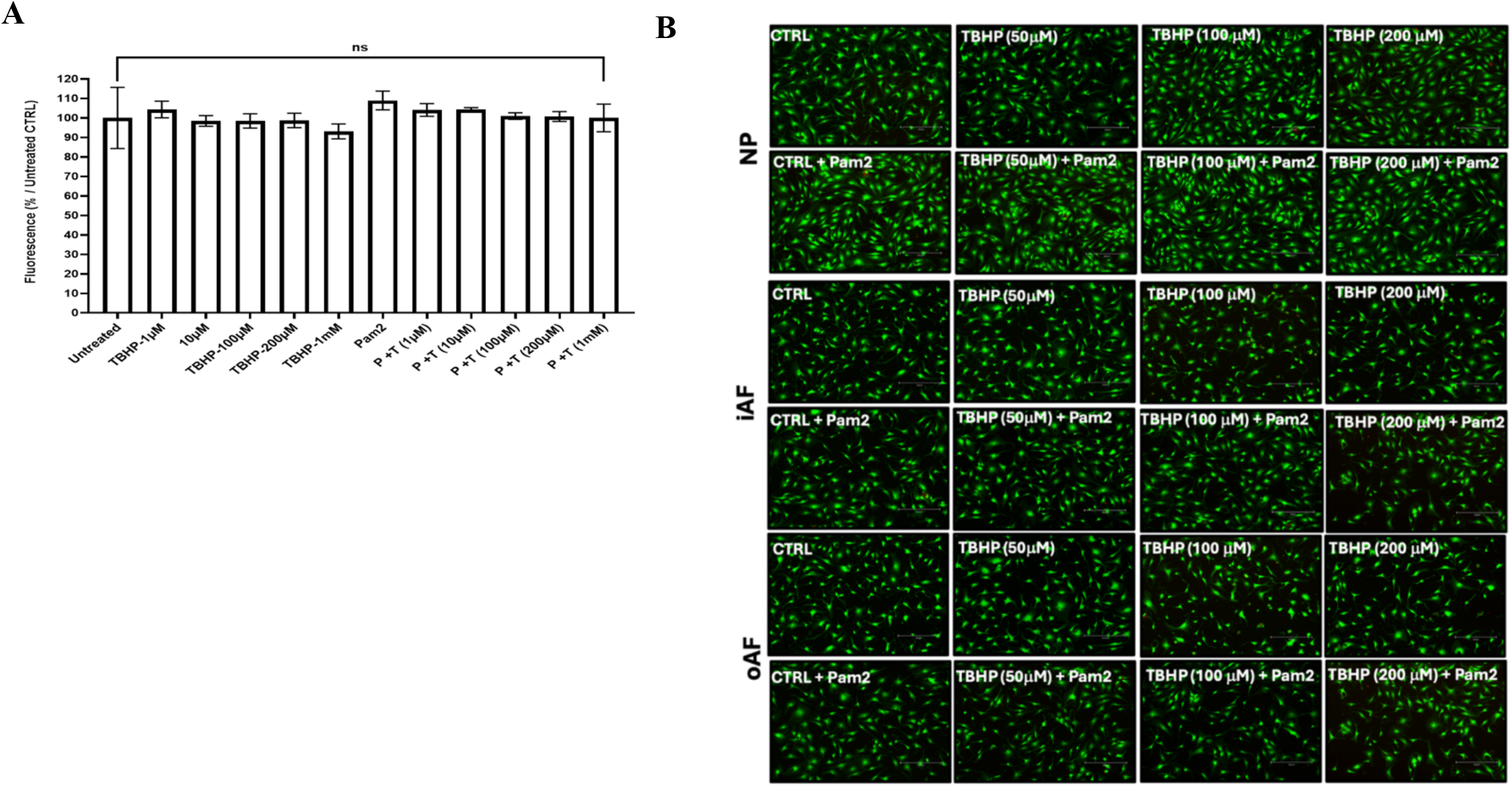
Cytotoxicity analysis of tBHP exposure. **A)** Metabolic activity of IVD cells following exposure to tBHP with and without Pam2CSK4 (100ng/ml). **B)** Cell viability across the three IVD cell populations: NP, iAF, and oAF, using the Live/Dead assay following exposure to tBHP at 50, 100, or 200 µM, either alone or in combination with Pam2CSK4 (100 ng/ml). Green fluorescence indicates viable cells and red fluorescence, dead cells. Data are presented as mean ± SEM, n = 3, assessed by repeated measures Analysis of Variance (ANOVA) with Turkey’s post hoc test for multiple pairwise comparison. Scale bars: 100 μm

### 2. Validation of senescence induction

#### 2.1. β-gal fluorescence intensity

After establishing that tBHP concentrations up to 1 mM, with or without TLR-2 activation, were non-cytotoxic, we next quantified senescence induction using the CellEvent Senescence Green Detection Kit (Thermo Fisher). Primary NP, iAF, and oAF cells were exposed to 100 ng/ml Pam2CSK4 followed by 25, 50, 100, 200, and 400 µM tBHP to determine the concentration that produced the strongest senescence induction. Across all three cell types, fluorescence intensity increased proportionally with tBHP concentration, indicating a dose-dependent response (Figure 2A). Among the tested doses, 200 µM tBHP produced the highest fluorescence intensity and was therefore selected for subsequent experiments. We then evaluated whether senescence induction at 200 µM tBHP was driven primarily by oxidative stress, inflammatory stimulation, or a combination. NP, iAF, and oAF cells were treated with Pam2CSK4 alone, tBHP alone, or Pam2CSK4 + tBHP, and fluorescence intensity was quantified relative to non-induced controls. Combined induction produced the strongest senescence response, increasing fluorescence intensity by 20% in NP (p > 0.05), 30% in iAF (p <0.001), and 50% in oAF cells (p < 0.0001) (**Figure 2B**). Together, these results demonstrate that oAF cells exhibit the strongest senescence response and combined induction, and that Pam2CSK4 + tBHP synergistically enhances senescence beyond either stimulus alone. Based on these findings, oAF cells treated with Pam2CSK4 (100 ng/ml) and 200 µM tBHP were selected in the subsequent experiments.

**Figure 2.**
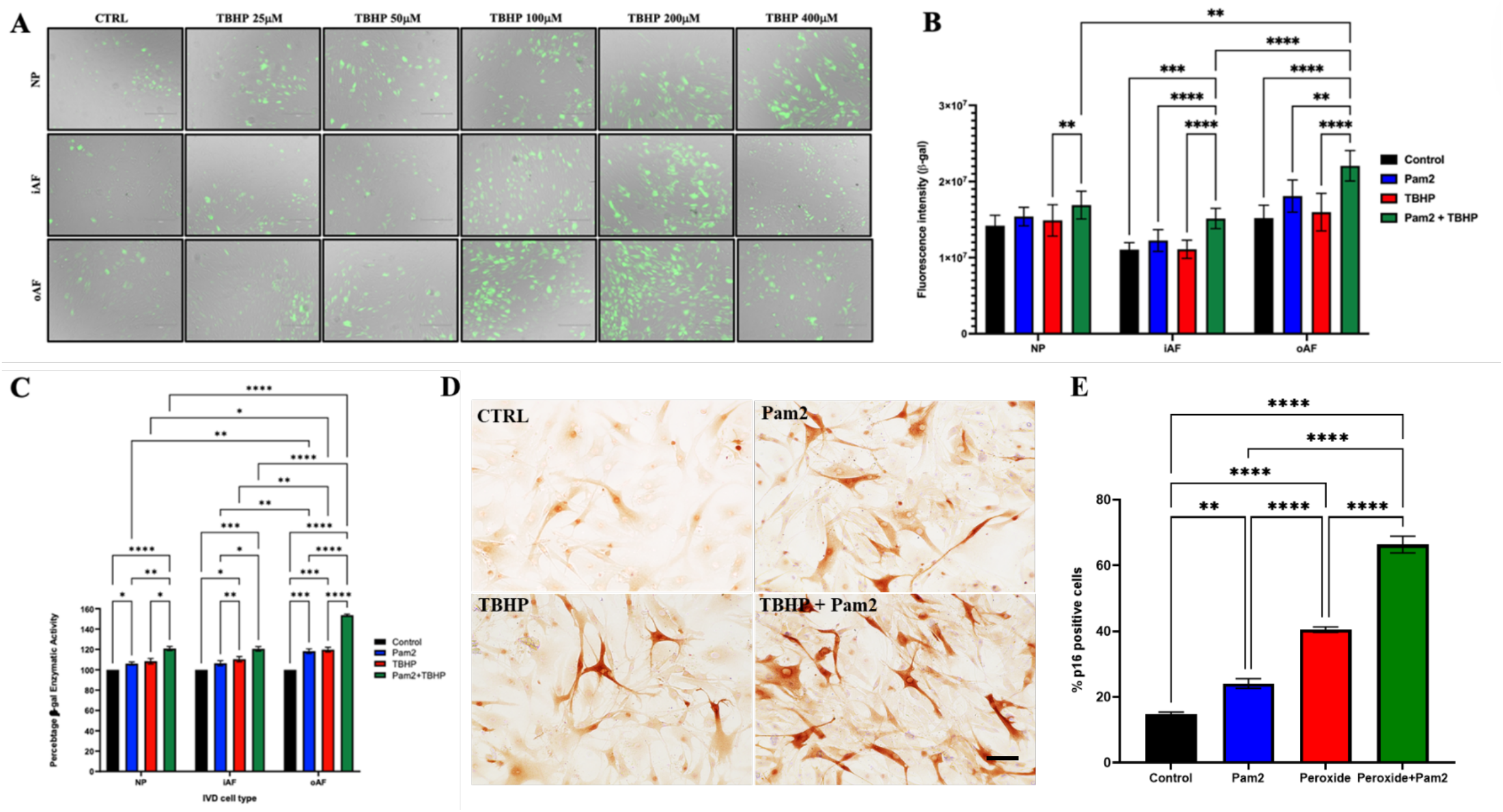
Measurement of senescence induction. **A)** Representative fluorescence micrographs showing SA-β-galactosidase fluorescence across nucleus pulposus (NP), inner annulus fibrosus (iAF), and outer annulus fibrosus (oAF) cells with and without senescence induction. Green fluorescence indicates β-gal-positive senescent cells overlaid on bright-field images. **B)** Quantification of β-gal fluorescence intensity **C)** Quantitative measure of β-gal enzymatic activity **D)** Evaluation of p16 expression in oAF. Brown nuclear staining indicates p16-positive senescent cells. **E)** Quantitative analysis of p16-positive cells. Data are presented as mean ± SEM, (n = 6), Scale bars: 100 μm in **(A)** and 50 μm in **(D).** *, **, ***, **** indicate significance of p ≤ 0.05, p ≤ 0.01, p ≤ 0.001, and p ≤ 0.001, respectively, assessed by repeated measures Analysis of Variance (ANOVA) with Turkey’s post hoc test for multiple pairwise comparison in **(B, C** and **E)**.

#### 2.2. β-gal enzymatic activity measurement

We next set out to evaluate whether findings obtained using the CellEvent Senescence Green Detection Kit were consistent with enzymatic β-galactosidase activity. For that, we used the Blue LacZ β-Gal Detection Kit (Abcam), a fluorometric assay that provides a sensitive and quantitative measure of senescence-associated β-galactosidase activity. Across the three IVD cell populations, β-gal activity following single and combined induction closely mirrored the fluorescence-based intensity results (**Figure 2B-C**). Combined induction with Pam2CSK4 + tBHP produced the strongest β-gal enzymatic activity. As with the intensity assay, single-agent induction produced a weaker increase. Consistent with the fluorescence-based intensity assay, oAF cells, under combined induction, exhibited the highest senescence response, showing a 55% (p < 0.0001) increase in β-gal activity, substantially greater than NP and iAF cells, which each showed about 20% increases (p ≤ 0.001) (**Figure 2C**). The agreement between assays confirms the robustness of the induction model and supports the use of Pam2CSK4 and tBHP as a model system for subsequent senolytic screening experiments.

#### 2.3. Confirmation of senescence using p16 as a marker

To determine whether β-galactosidase reflects true senescence rather than quiescence, we quantified expression of p16, a canonical marker of irreversible cycle arrest. Immunocytochemical staining of p16 in oAF cells following single-agent induction showed modest but significant increases relative to non-induced controls: Pam2CSK4 increased p16 expression by 10% (p ≤ 0.01), whereas tBHP increased p16 by 25% (p < 0.0001), These values were substantially lower than those observed with the combined induction. Consistent with the β-galactosidase results, combined Pam2CSK4 + tBHP induction demonstrated the strongest expression, increasing p16 by 55% (p < 0.0001) compared with untreated controls (**Figure 2D-E**). The similarity in response pattern observed for p16 expression and β-gal activity indicates that β-gal reflects the senescent cell levels and validates the robustness of the induction model for subsequent senolytic screening.

### B – Screening of toxicity and senolytic activity

#### 1. Measure of cytotoxicity and senolytic effects using metabolic activity

After establishing and validating the robustness and reproducibility of the senescence induction model, the next step was to establish a simple, low-cost screening method capable of reliably separating senolytic activity from general toxicity. Because senolysis is killing senescent cells, a reduction in the number of cells in induced cultures should correspond to a decrease in metabolic activity. Whereas non-induced cultures should be only minimally affected. We evaluated whether the Alamar blue metabolic activity method could serve as a screening method. We tested 3 well-characterized senolytics (o-vanillin, RG-7112, and ABT-199). First, we measured the metabolic activity in non-induced cells to confirm their non-toxic concentration range. Cells were exposed to increasing concentrations of o-vanillin (0.1-100 µM), RG-7112, and ABT-199 (0.01-100 µM), and metabolic activity was quantified. o-Vanillin and ABT-199 exhibited no detectable cytotoxicity, maintaining metabolic activity across all concentrations tested (**Figure 3A-C**). In contrast, RG-7112 was toxic at higher concentrations (100 µM) and reduced metabolic activity by 50% (Figure 3B). These results validated non-cytotoxic concentrations for the compounds and established the concentration range for subsequent validation of senolytic activity in senescence-induced cells (Cherif et al., 2019; Ji et al., 2016; Yosef et al., 2016; Colville et al. 2023).

**Figure 3.**
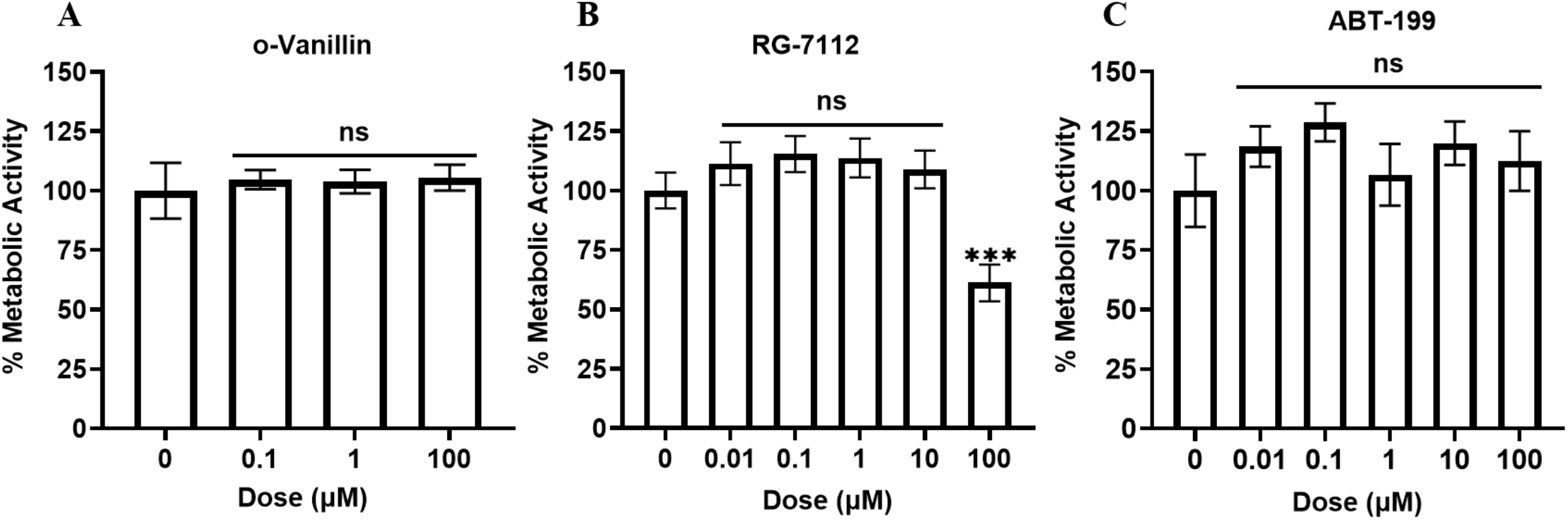
Metabolic activity measures established a non-cytotoxic range in non-induced cells. Metabolic activity was measured in non-induced cells following 6 h exposure to increasing concentrations of **(A)** o-vanillin (0.1-100 µM), **(B)** RG-7112 (0.01-100 µM), and ABT-199 (0.01-100 µM). Data are presented as mean ± SEM, n = 6. n.s and *** indicate no significance and significance of p ≤ 0.001, respectively, assessed by repeated measures Analysis of Variance (ANOVA) with Turkey’s post hoc test for multiple pairwise comparisons in **(A-C)**.

The senolytic effect of the compounds was then confirmed by a significant decrease in metabolic activity in senescence-induced cells in the non-toxic concentration range. o-Vanillin (0.1-100 µM) significantly reduced the metabolic activity by 20-30% (P ≤ 0.05), ABT-199 (1-100 µM) by 30-35% (P ≤ 0.05), and RG-7112 (0.1-10 µM) by 30-50% (P ≤ 0.05) relative to non-treated senescence-induced cells (**Figures 4A-C**).

**Figure 4.**
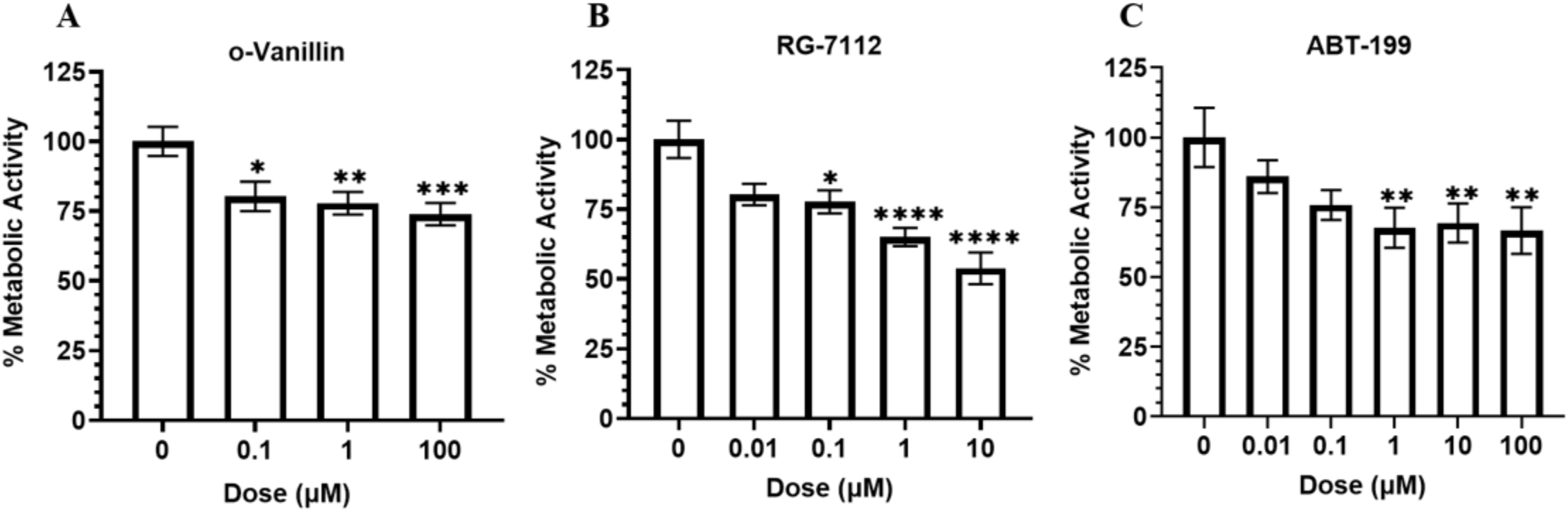
Metabolic activity in senescence-induced cells. **A)** Metabolic activity measure of induced oAF cells following 6 h exposure to o-vanillin (0.1-100 µM), **B)** RG-7112 (0.01-10 µM), and **C)** ABT-199 (0.01-100 µM). Data are presented as mean ± SEM, n = 6, *, **, ***, and **** indicate significance of p ≤ 0.05, p ≤ 0.01, p ≤ 0.001, and p ≤ 0.0001, respectively, assessed by repeated measures Analysis of Variance (ANOVA) with Turkey’s post hoc test for multiple pairwise comparison in **(A-C)**.

To validate that the reduction in metabolic activity truly reflected senolysis, we evaluated SA-β-gal enzymatic activity across the same senolytic concentration ranges. SA-β-gal activity decreased significantly in senescence-induced cells, confirming that the metabolic assay accurately reports killing of senescent cells. A consistent pattern emerged across agents: o-Vanillin reduced SA-β-gal activity by 20-30%. ABT-199 reduced SA-β-gal activity by 25-40%, and RG-7112 reduced SA-β-gal activity by 20-35%. All effects were calculated relative to vehicle-treated, senescence-induced controls (**Figure 5A-C**). These results confirm that metabolic activity can serve as a robust, low-cost, and sensitive readout for senolytic screening, capturing senolytic action across compounds with distinct potency profiles.

**Figure 5.**
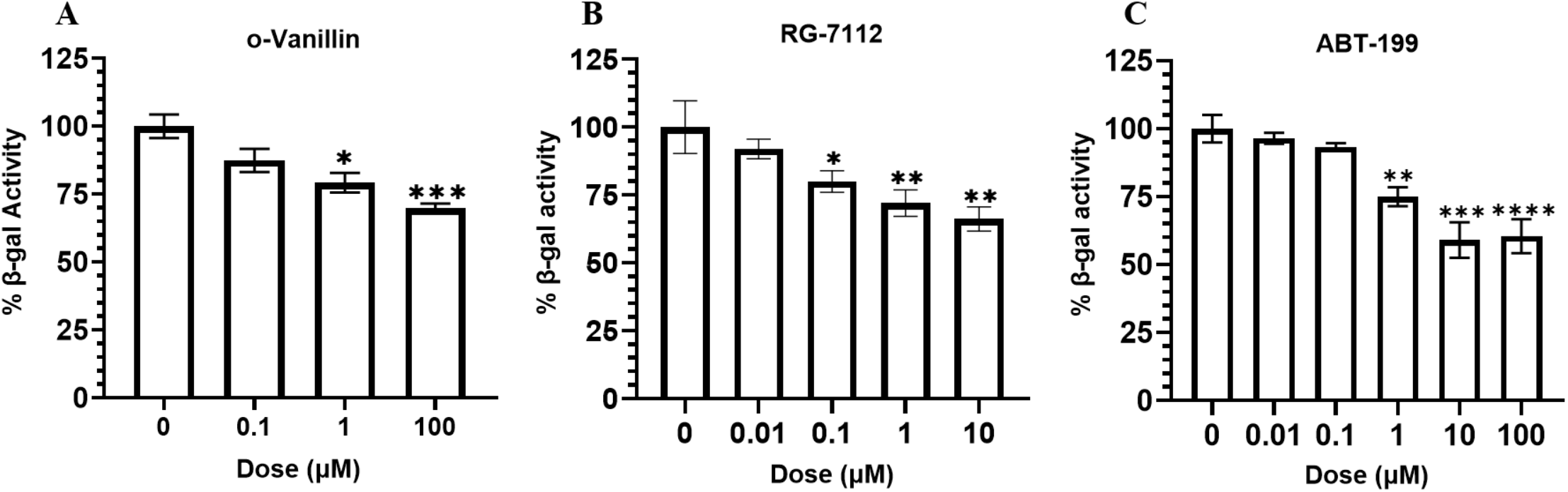
SA-β-gal enzymatic activity **validates the observed decrease in metabolic activity in induced cells.** SA-β-gal enzymatic activity of senescence-induced cells following 6 h exposure to increasing concentrations of **A)** o-vanillin, **B)** RG-7112, and **C)** ABT-199. Each compound reduced β-gal activity in a dose-dependent manner, consistent with the decrease in metabolic activity following the treatment. Data are presented as mean ± SEM, n = 6, *, **, ***, and **** indicate significance of p ≤ 0.05, p ≤ 0.01, p ≤ 0.001 and p ≤ 0.0001, respectively, assessed by repeated measures Analysis of Variance (ANOVA) with Turkey’s post hoc test for multiple pairwise comparison in **(A-C)**.

Finally, as an additional layer of validation, the expression of p16 was quantified at concentrations previously shown to have a senolytic effect. A significant decrease in p16-positive cells confirmed the SA-β-gal enzymatic activity assay results and validated the metabolic activity measure in induced cells as a screening method for senolytics (Figures 6A-B). o-Vanillin (100 µM), RG-7112 (5 µM), and ABT-199 (5 µM) significantly reduced the percentage of p16-positive cells to 17%, 25%, and 40%, respectively, representing a 75%, 60%, and 40% decrease, respectively, relative to vehicle-treated, senescence-induced controls. These results demonstrate that metabolic-activity measurements provide a functionally direct readout of senolysis, whereas p16 expression reflects pathway-specific phenotypic modulation.

**Figure 6.**
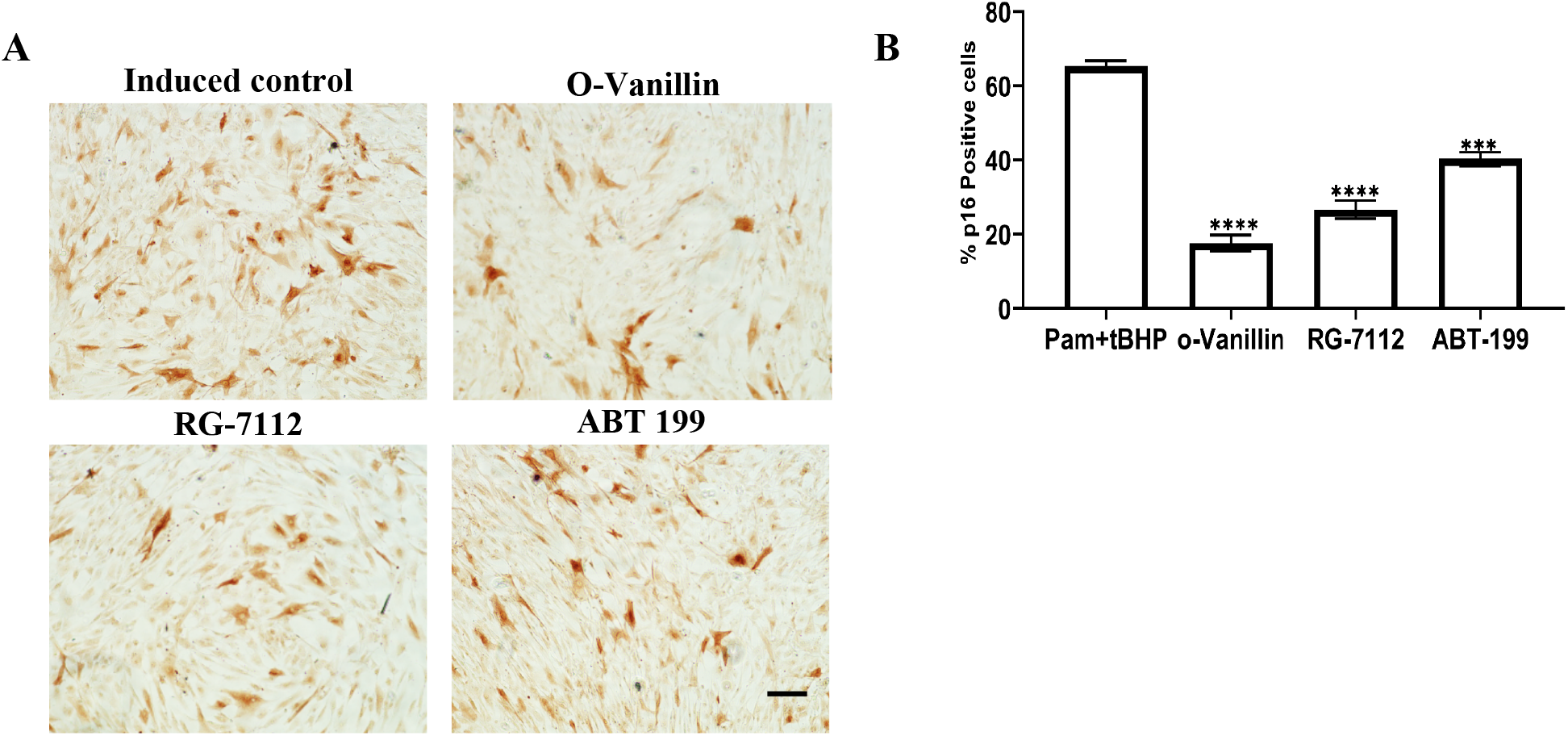
p16 expression after exposure to senolytics in induced cells. **A)** Representative immunohistochemical staining of senescence-induced oAF cells treated with senolytics: o-vanillin (100 µM), RG-7112 (5 µM), and ABT-199 (5 µM). Brown nuclear staining indicates senescent cells. **B)** Quantitative analysis of p16-positive cells. Data are presented as mean ± SEM, n = 6, Scale bars: 100 μm in **(A).** *** and **** indicate significance at p ≤ 0.001, and p ≤ 0.0001, respectively, assessed by repeated-measures Analysis of Variance (ANOVA) with Tukey’s post hoc test for multiple pairwise comparisons in **(B)**.

Our findings demonstrate that the presented induction method generates a standardized and treatment-responsive senescent platform and that metabolic activity measurements provide a simple, low-cost, sensitive, rapid, and practical primary screening assay for identifying compounds with selective senolytic activity. Validation using three mechanistically distinct senotherapeutics (o-Vanillin, RG-7112, and ABT-199) and orthogonal senescence readouts, including SA-β-gal activity and p16 immunostaining, supports the robustness and reproducibility of this platform for screening and prioritizing candidate senolytic compounds.

## Discussion

In this study, we established and validated a standardized in vitro method for senescence induction and senolytic drug screening using primary human IVD cells as a representative model. Many existing senescence induction models rely on replicative exhaustion, genotoxic stress, or irradiation, which do not recapitulate the biological drivers of senescence in musculoskeletal tissues (De Magalhães et Passos, 2018; Gresham et al., 2022; Ou et al., 2021; Veronesi et al., 2023). Degenerating musculoskeletal tissues experience chronic exposure to inflammatory cytokines (Le Maitre 2005; Risbud & Shapiro 2014; Phillips 2013; Sun et al., 2018; Wang et al., 2011) and oxidative stress arising from impaired nutrient transport, mitochondrial dysfunction, and mechanical overload (Vergroesen et al., 2015; Hou et al., 2025; Chen et al., 2024). By combining inflammatory stress using TLR-2 activation with Pam2CSK4 (Mannarino et al., 2021; Krock et al., 2016, 2017) and oxidative stress induced by tBHP (Qiu et al., 2024; Yang et al., 2021), we generated a senescent cell population that mimics the physiological inflammatory-oxidative microenvironment characteristic of degenerating human MSK tissues (Unterluggauer, 2003; Yang et al., 2014; Yeh et al., 2020). This dual-stimulus model produced a robust and reproducible senescence response across NP, iAF, and oAF cells. Across all assays, oAF cells demonstrated the highest senescence induction. This observation aligns with our previous report of a higher proportion of p16^*INK4a*^-positive cells in AF compared to NP tissue (Cherif et al., 2019). Moreover, previous reports have shown that annulus fibrosus cells exhibit greater susceptibility to mechanical and oxidative stress. The oAF region experiences the greatest tensile loading during spinal motion (Ning et al., 2021; Dai et al., 2022).

Our combined induction model was validated by a stronger increase in three independent senescence markers, including β-gal fluorescence, β-gal enzymatic activity, and p16 expression (Gorgoulis et al., 2019; Hernandez-Segura et al., 2018). The concordance across these complementary assays provides strong evidence that the induction model generates true senescence rather than transient quiescence. Although p16 immunostaining, SA-β-gal enzymatic activity, and metabolic activity exhibited consistent directional trends across both induction and senolytic treatment conditions, differences in effect size are expected because these assays quantify distinct biological hallmarks of senescence (González-Gualda et al., 2021). p16 immunostaining quantifies the proportion of p16-positive cells under irreversible cell-cycle arrest, SA-β-gal enzymatic activity reflects lysosomal expansion at the population level, and metabolic assays measure the overall reducing capacity of viable cells. Because these assays quantify distinct biological features of senescence using different measurement principles, differences in effect size are expected even when they demonstrate the same overall directional response. Such assay-specific differences are widely reported and explain why p16, SA-β-gal activity, and metabolic activity frequently correlate in direction but differ somewhat in magnitude (González-Gualda et al., 2021). In our results, the ranking of senolytic potency based on p16 differed from that observed with metabolic activity and SA-β-gal enzymatic activity, with o-vanillin producing the strongest reduction in p16 expression. This divergence most likely reflects the distinct biological endpoints measured by each assay and may also be influenced by the different mechanisms of action of the senolytic compounds. We previously reported that o-vanillin modulates NRF2- and NF-κB-associated signalling, whereas RG-7112 and ABT-199 primarily promote senescent cell apoptosis through p53 stabilization and BCL-2 inhibition, respectively (Cherif et al., 2019, 2020; Mannarino et al., 2020, 2021; Yosef et al., 2016).

Metabolic activity is not a senescence-specific marker but rather a widely used measure of cellular metabolic function and viability (O’Brien et al., 2000; Rampersad, 2012). Despite the permanent state of cell growth arrest, senescent cells are highly metabolically active and fuel a massive output of proteins, lipids, and inflammatory signals (Wiley et Campisi, 2016). Thus, we used metabolic activity measurement as a readout to distinguish senolytic compound activity in induced senescent cells from general cytotoxicity in non-induced non-senescent cells. We validated our approach using three known senolytics, o-Vanillin, RG-7112, and ABT-199, at different concentration ranges (Cherif et al., 2019, 2020; Mannarino et al., 2020, 2021; Yosef et al., 2016; Colville et al., 2023).

We determined a window where the three senolytics were non-toxic in non-senescent cell populations but significantly reduced metabolic activity in senescence-induced cells, demonstrating their selective senolytic activity. This decrease in metabolic activity reflects the elimination of metabolically active senescent cells and confirms the utility of metabolic activity as a primary screening readout. Importantly, these reductions correlated with decreases in SA-β-gal enzymatic activity and nuclear p16 accumulation, validating that metabolic activity accurately reports senolysis. The senolytic profiles observed for o-vanillin, RG-7112, and ABT-199 are consistent with previously reported activities for these compounds (Cherif et al., 2019, 2020; Mannarino et al., 2020, 2021; Yosef et al., 2016; Colville et al., 2023).

The metabolic activity assay is described here as a senolytic screening platform that can be readily integrated with other senescence-induction strategies depending on the biological question or clinical context being modelled. Although Pam2CSK4+tBHP induction captures the inflammatory-oxidative environment characteristic of intervertebral disc degeneration, alternative induction methods, such as radiation-induced, chemotherapy-induced, or replicative senescence, may be more appropriate for senotherapeutics intended for oncology, hematology, or systemic aging indications. This alignment is essential because one of the major causes of senolytic trial failure may have been the mismatch between preclinical models and the senescent cell populations present in human disease (Lane et al., 2021; Nambiar et al., 2023). Many early-stage assays rely on senescent fibroblasts or immortalized cell lines, which do not recapitulate the phenotype, stress-response pathways, or drug vulnerabilities of senescent cells in vivo. When senolytics are evaluated in models that do not reflect the biology of the target tissue, such as the disc, lung, kidney, or cartilage, the resulting efficacy signals often fail to translate clinically. Generating senescent cell populations that mirror the disease-specific senescence phenotype therefore increases the predictive value of early-stage screening and strengthens the translational potential of senolytic candidates ^14^.

## Conclusion

By integrating TLR-2-mediated inflammatory stimulation with oxidative stress, the induction model incorporates two major pathological drivers of naturally occurring cell senescence in MSK tissues. This induction method generates an increase in senescent cells validated using multiple independent senescence markers. We further demonstrate that a metabolic activity assay can distinguish senolytic activity from general cytotoxicity when interpreted within our standardized senescence induction protocol. Screening results, using metabolic activity, correlate with reductions in SA-β-gal activity and p16 expression, as validated by three distinct senolytics. Collectively, these findings support the use of the induction protocol and the senolytic screening method as a biologically and clinically relevant method for the primary screening of senolytic compounds.

## Material and methods

### Schematic representation of the experimental workflow

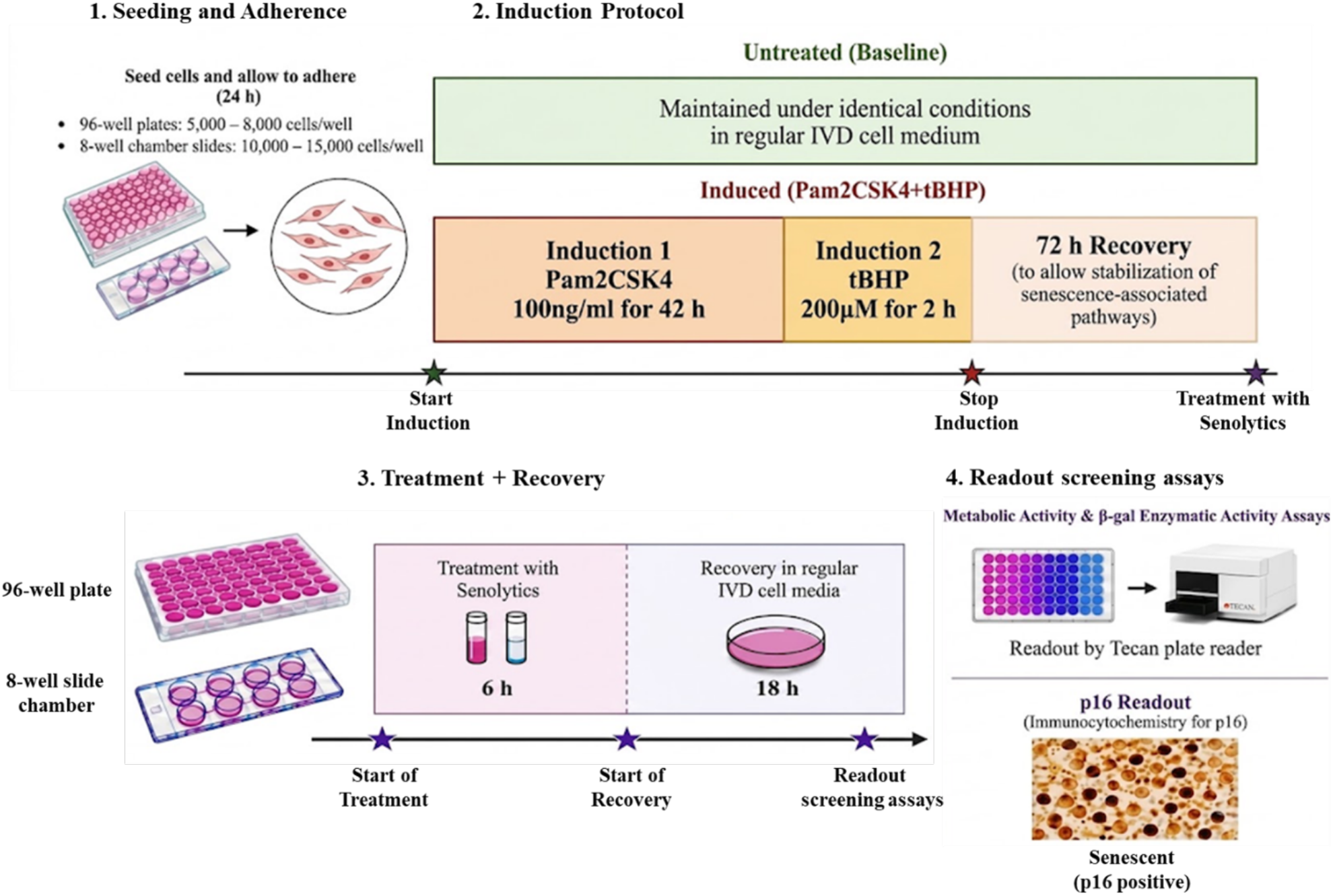

### Tissue Collection and Cell Isolation

Briefly, human lumbar IVDs were retrieved from spines obtained with familial consent (IRB00010120) through the Transplant Quebec Organ Donation Program. Discs dissected from the spinal column were used for cell isolation. Nucleus pulposus (NP), inner annulus fibrosus (iAF), and outer annulus fibrosus (oAF) cell populations were isolated from discs as previously described ^15^. Samples were washed in phosphate-buffered saline solution (Sigma-Aldrich, Oakville, ON, Canada) and Hank’s-buffered saline solution (HBSS, Sigma-Aldrich, Oakville, ON, Canada) supplemented with PrimocinTM (InvivoGen, San Diego, CA, USA). Then, the tissue was minced and digested in 0.15% collagenase type II (Gibco) for 16 h at 37 °C. Cells were passed through both a 100 μm filter and a 70 μm filter before being resuspended in Dulbecco’s Modified Eagle Medium (DMEM, Sigma-Aldrich, Oakville, ON, Canada) supplemented with 10% fetal bovine serum (FBS, Gibco), Primocin™, and Glutamax (Oakville, ON, Canada) and maintained in a 5% CO_2_ incubator at 37°C. Cells were used between passages 1 and 2 to preserve phenotype.

### Senescence induction

Cells were seeded at a density of 8-10K cells/well in 96-well plates/10-15K/chamber in 8-well chamber slides and allowed to adhere for 24 h before induction. Senescence was induced using single induction or a combined sequential induction protocol. Cells were exposed to Pam2CSK4 alone at 100 ng/mL for 42 h, tBHP alone at 200 µM for 2 h, or combined induction, consisting of Pam2CSK4 (100 ng/mL) for 42 h, followed by tBHP (200 µM) for 2 h, and subsequently a 72-h recovery period to allow stabilization of senescence phenotype. Non-induced cultures maintained under identical conditions served as controls. The 200 µM tBHP concentration was pre-validated to be non-cytotoxic while still eliciting a robust senescence phenotype, ensuring that downstream readouts reflected senescence rather than cell death. The oAF cell population that exhibited the strongest senescence response was selected to conduct downstream experiments.

### Treatment with senolytic agents

Non-induced oAF cells were treated in 96-well plates for 6 h with three known senolytics, followed by 18 h of treatment recovery. The cells were exposed to a range of concentrations of o-Vanillin (0.1-100 µM), RG-7112 and ABT-199 (0.01-100 µM) to evaluate the cytotoxic profile in non-senescent oAF cells. To evaluate the senolytic selectivity, non-cytotoxic ranges of concentrations were tested in a senescence-induced cell population.

### Cell viability assay

Cells were seeded in 96-well plates, and viability was assessed using the LIVE/DEAD™ Viability/Cytotoxicity Kit for Mammalian Cells (Thermo Fisher Scientific, Mississauga, ON, Canada) according to the manufacturer’s instructions. Briefly, culture medium was removed, cells were washed once with PBS, and 150 µL of the freshly prepared LIVE/DEAD working solution (containing ~2 µM calcein-AM and ~4 µM EthD-1) was added to each well. Cells were incubated for 45 min at room temperature, protected from light, and fluorescence was imaged immediately to identify viable (green, calcein-positive) and non-viable (red, EthD-1–positive) cells.

### Metabolic activity

Metabolic activity was quantified using an Alamar Blue assay as previously described ^13, 16^. Briefly, Cells were exposed to 10% Alamar Blue reagent (Thermo Fisher, Waltham, MA, USA) in DMEM and incubated for 2-4 h at 37 °C. Fluorescence (Ex560/Em590) was measured using a spectrophotometer (Tecan Infinite T200, Männedorf, Switzerland) equipped with Magellan software (Tecan, Männedorf, Switzerland). The results are presented as a percentage of metabolic activity compared to the control. All experiments were performed with n = 6 biological replicates, each measured in duplicate wells per condition.

### SA-β-Gal Staining

SA-β-gal staining was carried out on cells that were seeded in 8-well chamber slides (Thermo Fisher Scientific, Mississauga, ON, Canada), according to the manufacturer’s protocol (Thermo Fisher Scientific, Mississauga, ON, Canada). Following treatment, the culture media was removed, and the cells were washed twice with PBS, then fixed with the provided fixation buffer for 10 min at room temperature. After rinsing three times with PBS, the staining mixture was added, and the plate was sealed with Parafilm and incubated for 2 h at 37 °C in an incubator without CO_2_. Chamber slides were mounted with Aqua Polymount (Polysciences, Warrington, PA, USA) and visualized with the EVOS™ M5000 Imaging System (Thermo Fisher Scientific, Waltham, MA, USA). Ten fields, randomly distributed across the well, were analyzed, and the fluorescence intensity was measured using QuPath bioimage analysis (Bankhead et al., 2017). All experiments were performed with n = 6 biological replicates, each measured in duplicate.

### SA-β-gal Enzymatic Activity

SA-β-galactosidase enzymatic activity was quantified using the Abcam FACS Blue LacZ kit (Abcam, Cambridge, Ma, USA) as described ^13^. Briefly, culture medium was removed, and cells were washed with PBS (Sigma-Aldrich, Oakville, ON, Canada). Cells were then incubated with 50 µL/well of Reaction Buffer (200 mM sodium phosphate buffer, pH 7.0; 100 mM MgCl_2_; 1 M β-mercaptoethanol; 10% Triton X-100) for 5 min at room temperature. Subsequently, 50 µL/well of LacZ fluorogenic substrate was added, and cells were incubated for 20 min. The reaction was stopped by adding 50 µL/well of Stop Buffer (500 mM glycine and 10 mM EDTA in ddH_2_O, pH 12). Fluorescence was measured at Ex 390 nm / Em 460 nm using a Tecan Infinite T200 spectrophotometer (Männedorf, Switzerland) equipped with Magellan software. β-galactosidase activity was expressed as a percentage relative to the control group. All experiments were performed with n = 6 biological replicates, each measured in duplicate.

### P16 immunocytochemistry staining

Cells were seeded in 8-well chamber slides, and p16^**INK4a**^ immunocytochemistry staining was performed on monolayer culture using the P16 antibody (CINTec Kit, Roche) as described before ^13, 16^. Briefly, cells were fixed with 4% PFA, washed three times with PBS, and blocked with 1% BSA, 1% goat serum, and 0.1% Triton X-100 for 30 min. All samples were incubated at 4 °C overnight with p16INK4a antibody and PBS-T for negative control. Secondary detection steps were performed according to the manufacturer’s instructions. Afterwards, Chamber slides were mounted with Aqua Polymount (Polysciences, Warrington, PA, USA). Images were captured with EVOS™ M5000 Imaging System (Thermo Fisher Scientific, Waltham, MA, USA). Ten fields, randomly distributed across the well, were analyzed, and the fluorescence intensity was measured using QuPath bioimage analysis (Bankhead et al., 2017). The results are presented as a percentage of p16-positive cells. All experiments were performed with n = 6 biological replicates, each measured in duplicate.

### Statistical analysis

Data were analyzed using GraphPad Prism 10.6.1 (GraphPad Software, La Jolla, CA, USA). Because all experiments involved matched samples from the same donors across multiple treatments, analyses were performed using repeated-measures ANOVA, followed by Tukey’s post hoc test for multiple pairwise comparisons. The specific statistical test and correction applied for each experiment are indicated in the corresponding figure legends. A p-value < 0.05 was considered statistically significant. All data are presented as mean ± SD.

## Notes

### Competing Interest Statement

The authors have declared no competing interest.

### Funding

- Canadian Institutes of Health Research: PJT-178111
- Arthritis Society: SOG-20-0000000075

